# Right atrial mechanism contributes to atrial fibrillation in a canine model of pulmonary arterial hypertension

**DOI:** 10.1101/2022.02.19.481116

**Authors:** Yang Liu, Xiyao Zhu, Ziliang Song, Mu Qin, Changhao Xu, Xu Liu

## Abstract

**Objectives:** The aim of this study was to investigate proarrhythmic substrates of atrial fibrillation (AF) in a canine model of dehydromonophylline (DMCT)-induced pulmonary arterial hypertension (PAH).

**Methods:** All cannines (n=12) underwent baseline echocardiographic and hemodynamic examinations, 7 of which were injected with DMCT (3.0mg/kg) to induce PAH via jugular vein cannulation. The control beagles (n=5) were given the same dose of normal saline. Then, both groups were monitored by insertable cardiac monitors. Hemodynamic, echocardiographic, electrophysiological and histological examinations were performed 8 weeks later.

**Results:** In PAH group, 2 died after the injection (mortality 28.6%). Thus, 10 beagles (PAH group: 5, control group: 5) underwent all the examinations. The pulmonary artery pressure increased significantly while the right atrium (RA) and right ventricle expanded slightly. Spontaneous AF episodes were recorded in all PAH canines 1 week after the injection. The AF burden increased rapidly from 1 week (7.6±1.8%) and remained high after 2-3 weeks (32.0±4.9% at 8 weeks). Compared with the control group, the PAH group had abbreviated effective refractory periods (ERPs), increased atrial ERP dispersion, and slower conduction velocities. Notably, AF susceptibility and atrial remodeling in RA was more significant those in LA, such as increased WOV(39.0±6.5ms vs. 28.0±5.7ms, P=0.022), enlarged low voltage regions (7.66±0.46% vs. 4.40±0.55%, P<0.0001) and fibrosis (8.22±0.61% vs. 4.93±0.60%, P <0.0001).

**Conclusions:** DMCT-induced canine PAH model increased the incidence of spontaneous and induced AF. The electrophysiological and structural remodeling of the RA facilitated the AF genesis.

## Introduction

Pulmonary arterial hypertension (PAH) is associated with significant morbidity.^[1-3]^ In recent years, multicenter studies have confirmed that elderly patients have accounted for more than half of the total PAH population.^[4]^ A United States population-based survey also found that hospitalization rates for PAH have increased over the past 20 years, and that 30% of the patients dying with PAH were 75 years of age or older.^[5]^ Notably, patients with PAH often have atrial fibrillation (AF),^[6]^ and persistent AF was associated with a 2-year mortality>80%.^[7]^ However, because of the lack of large animal models of AF due to PAH, the exact mechanism of PAH leading to AF is still unclear. Therefore, it is impossible to formulate corresponding intervention strategies. Previous study demonstrated that right atrium (RA) can be remodeled by longstanding pressure and volume overload in PAH rat model, seemingly producing the underlying arrhythmogenic substrate.^[8]^ However, due to the lack of studies on large animal models of PAH induced AF, the triggering and maintenance mechanisms of AF remain unclear. The purpose of this study was to explore the the alteration of pro-arrhythmic substrate in PAH induced AF canine model.

### Methods

All protocols conformed to the Guide for the Care and Use of Laboratory Animals published by the US National Institutes of Health (NIH Publication No. 85–23, revised 1996). Experiments were approved by the Animal Care and Use Committee of the Shanghai Chest Hospital, Shanghai Jiao Tong University.

### Animal preparations

Dehydromonocrotaline (DMCT) was prepared as previously described^[9]^ and was dissolved in dimethylformamide (0.1ml/kg). Procedures were performed on healthy adult male beagles (21.0±1.5kg) that were anesthetized (25 mg/kg intravenous [IV] pentobarbital sodium) and ventilated with room air with a positive pressure respirator (Harvard Apparatus Co., Natick, MA, USA). In the PAH group, an 11F hemostasis introducer (Abbott Medical, MN, USA) was placed in the right jugular vein and a diagnostic ultrasound catheter (Biosense Webster, CA, USA) was passed through the hemostasis introducer into the right cardiac system for echocardiographic examinations. Then, a 5-F Swan-Ganz catheter (Cordis, Miami, FL) was advanced through the 11F hemostasis introducer into the RA, the right ventricle (RV) and the pulmonary artery (PA). Hemodynamic measurements were performed in lateral position. Continuous hemodynamic monitoring included heart rate, pulmonary artery pressure (PAP), right ventricular pressure (VRP) and transcutaneous oxygen saturation. DMCT (3.0mg/kg) was administered via the main pulmonary artery. Then, an insertable cardiac monitor (ICM) (St Jude Medical, MN, USA) was implanted under the chest skin of the beagle to monitor for arrhythmia events (**Figure 1**). In the control group, the beagles underwent echocardiographic examinations, hemodynamic examinations and implanted ICM without DMCT injection. 8 weeks later, hemodynamic examinations, echocardiographic examinations, electrophysiological examinations and histopathological studies were performed.

**Figure 1.**
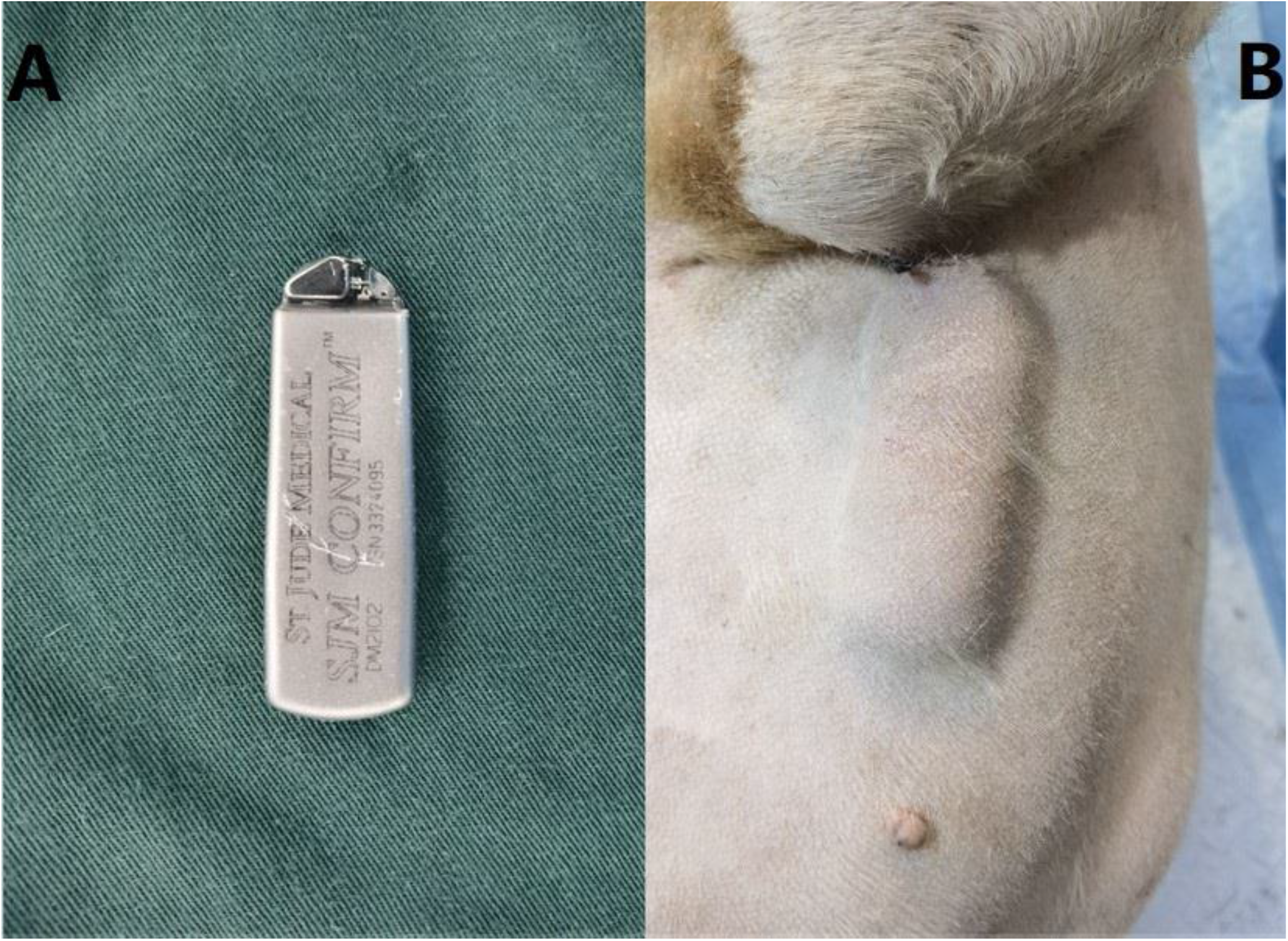
Insertable cardiac monitor (A) and its subcutaneous condition (B).

### Hemodynamic and echocardiographic examinations

8 weeks later, a 5-F Swan-Ganz catheter and a diagnostic ultrasound catheter were respectively delivered into the right cardiac system through an 11F hemostasis introducer placed in the left jugular vein, and the hemodynamic and echocardiographic examinations were performed successively according to the methods described above.

### Electrophysiological examinations

#### Electroanatomical mapping

A PentaRay catheter (Biosense Webster, CA, USA) was inserted into RA through the 11F hemostasis introducer placed in the left jugular vein, and the endocardial electroanatomical mapping of RA was performed under the guidance of the Carto 3 system (Biosense Webster). Then, the PentaRay catheter was placed into RV for endocardial electroanatomical mapping. After that, the chest was opened using a left thoracotomy and the pericardium was opened. The PentaRay catheter was placed into the pericardium via incision for epicardial electroanatomical mapping. Finally, an 8F hemostasis introducer (St Jude Medical, MN, USA) was inserted at the apex of left ventricle (LV), and the PentaRay catheter was sent into the left cardiac system through the 8F hemostasis introducer for the endocardial electroanatomical mapping of left atrium (LA) and LV.

#### Programmed stimulation

A programmed electrical stimulation (PES) protocol was used to examine the atrial and ventricular effective refractory period (ERP) and arrhythmia inducibility. Briefly, PES consisted of an eight-stimulus (S1) drive train followed by a ninth extra stimulus (S2). The CL of the S1 train was 500ms. The S1–S2 interval started at 350ms and was progressively reduced by 5ms per cycle until the S2 stimulus could no longer evoke atrial or ventricular deflection. ERP was defined as the longest S1–S2 interval that caused loss of atrial or ventricular depolarization. We measured the ERP of different regions of the atrium and ventricle, and used dispersion of ERP (dERP) to represent the differences in ERP among different regions of the atrium and ventricle. The difference between the maximum ERP and the minimum ERP divided by the mean of the ERP was the value of dERP. During ERP measurements, if AF was induced by decremental S1-S2 stimulation, the longest minus the shortest S1-S2 interval at which AF was induced was designated as the width of the window of vulnerability (WOV), which was used to evaluate the inducibility of AF.

#### Myocardial conduction mapping

The conduction time of the LA and RA were calculated by using the bipolar and unipolar potential information collected under sinus rhythm and the LAT software module and isochronal function of CARTO 3 system. Then, the conduction distance of myocardium was measured by the design line function. Thus the conduction velocity of myocardium could be calculated.

### Histopathological examinations

After the above examinations, tissue preparations (1×1.5cm) were separated from the atrial tissues and stored. Masson’s trichrome staining for identifying collagen fibers was used in the atrial preparations. Images captured for each preparation were analyzed using Image-pro plus 6.0 (Media Cybernetics, Inc., Rockville, MD, USA) to quantify the degree of fibrosis.

### Statistical analysis

All continuous variables were expressed as mean ± SD. Unpaired Student’s t test was used to evaluate the differences between baseline and 8-week values and between the PAH and control group. A P value of < 0.05 was considered to be statistically significant. SPSS version 19.0 and GraphPad Prism version 8.2.1 were applied.

## Results

7 beagles were injected with the DMCT, and 2 died suddenly during the injection (mortality 28.6%). 5 beagles were included in the control group. Thus, 10 beagles (PAH group: 5, control group: 5) underwent the following series of experiments.

### Hemodynamic and echocardiographic examinations

Compared with their baseline values, pulmonary arterial systolic pressure (PASP), pulmonary artery mean pressure (PAMP), right ventricular systolic pressure (RVSP), right ventricular mean pressure (RVMP) were significantly higher after 8 weeks in the PAH group (PASP: 35.6±3. 7mmHg vs. 19.0±2. 2mmHg, P<0.0001; PAMP: 23.8±3.2 mmHg vs. 11.0±2. 1mmHg, P<0.0001; RVSP: 33.4±3. 2mmHg vs. 17.4±2. 3mmHg, P< 0.0001; RVMP: 18.4±3.0mmHg vs. 8.6±2. 1mmHg, P<0.0001). Compared with the baseline values, the pulmonary arterial dimension (PAD) of the PAH beagles were significantly higher (PAD: 16.5±0.4mm vs. 15.1±0.4mm, P =0.001), while the right atrial dimension (RAD) and right ventricular diastolic dimension (RVD) were only slightly higher (RAD: 30.2±1. 2mm vs. 28.8±1.2mm, P=0. 127; RVD: 24.1±1. 5mm vs. 22.6±1.0mm, P=0. 120). In addition, the PAH group had mild to moderate pulmonary valve regurgitation and tricuspid regurgitation, which were absent at baseline (**Figure 2**). In the control group, none of these above items changed significantly from baseline after 8 weeks. The above results were shown in **Table 1**.

**Table 1.** Changes in echocardiographic parameters and hemodynamic parameters at baseline and after 8 weeks in PAH group and control group

| Characteristics | PAH group |  | Control group |  |
| --- | --- | --- | --- | --- |
|  | Baseline | After 8 weeks | Baseline | After 8 weeks |
| Echocardiographic parameters |  |  |  |  |
| RAD (mm) | 28.8±1.2 | 30.2±1.2 | 29.6±1.3 | 29.5±1.8 |
| LAD (mm) | 27.2±0.9 | 27.3±0.8 | 27.9±1.0 | 28.0±1.3 |
| RVDD (mm) | 22.6±1.0 | 24.1±1.5 | 23.6±1.0 | 23.8±1.3 |
| LVDD (mm) | 36.8±1.5 | 36.7±1.1 | 37.5±1.1 | 37.7±1.5 |
| PAD (mm) | 15.1±0.4 | 16.5±0.4* | 15.1±0.6 | 15.0±0.7 |
| LVEF (%) | 66.0±3.3 | 66.5±1.5 | 66.0±3.6 | 66.7±3.2 |
| Hemodynamic parameters |  |  |  |  |
| PASP (mmHg) | 19.0±2.2 | 35.6±3.7* | 19.8±2.4 | 19.2±2.4 |
| PAMP (mmHg) | 11.0±2.1 | 23.8±3.2* | 12.4±2.3 | 11.2±1.9 |
| RVSP (mmHg) | 17.4±2.3 | 33.4±3.2* | 18.2±1.9 | 17.6±2.1 |
| RVMP (mmHg) | 8.6±2.1 | 18.4±3.0* | 8.8±2.4 | 10.2±1.6 |

PAH: pulmonary arterial hypertension; RAD: right atrial dimension; LAD: left atrial dimension; RVDD: right ventricular diastolic dimension; LVDD: left ventricular diastolic dimension; PAD: pulmonary arterial dimension; LVEF: left ventricular ejection fraction; PASP: pulmonary arterial systolic pressure; PAMP: pulmonary arterial mean pressure; RVSP: right ventricular systolic pressure; RVMP: right ventricular mean pressure.
Data are presented as the mean ±SD. \*P<0.05 compared with the baseline.

**Figure 2.**
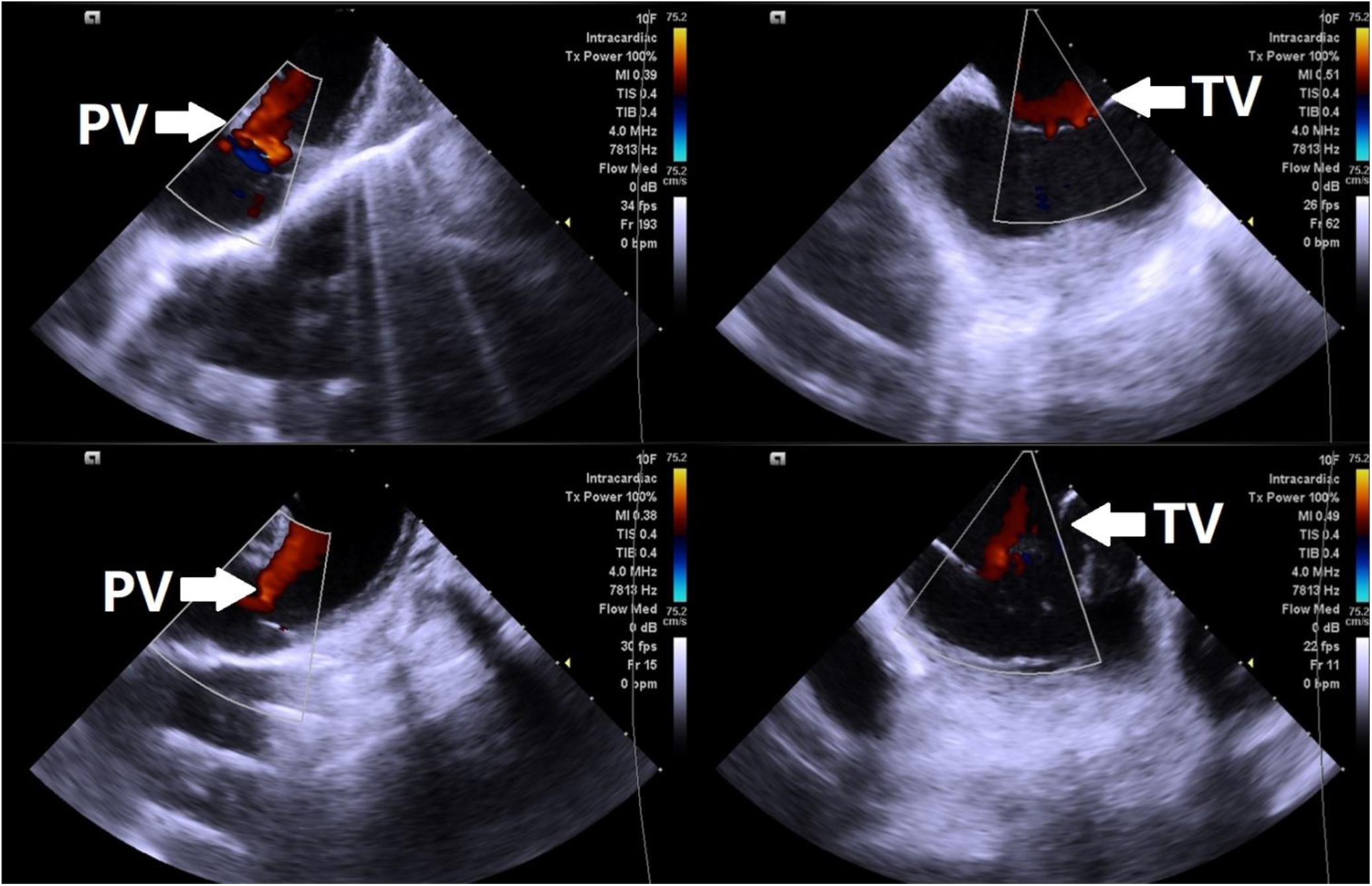
Pulmonary valve regurgitation and tricuspid regurgitation in the PAH group 8 weeks after the injection of dehydromonocrotaline. PV: pulmonary valve; TV: tricuspid valve.

### Arrhythmogenesis and electrophysiological examinations

#### Results of ICM

according to the data recorded by ICM, AF was recorded in all the PAH beagles 1 week after the injection of DMCT, and the AF burden was 7.6±1.8% at 1 week. The AF burden increased rapidly 2-3 weeks after the injection of DMCT (31.0±5.4% at 3 weeks). Then, the AF burden remained high, and the value was 32.0±4.9% at 8 weeks. (**Figure 3)** Ventricular arrhythmias were not recorded in the PAH beagles. In the control group, atrial and ventricular arrhythmias were not recorded.

**Figure 3.**
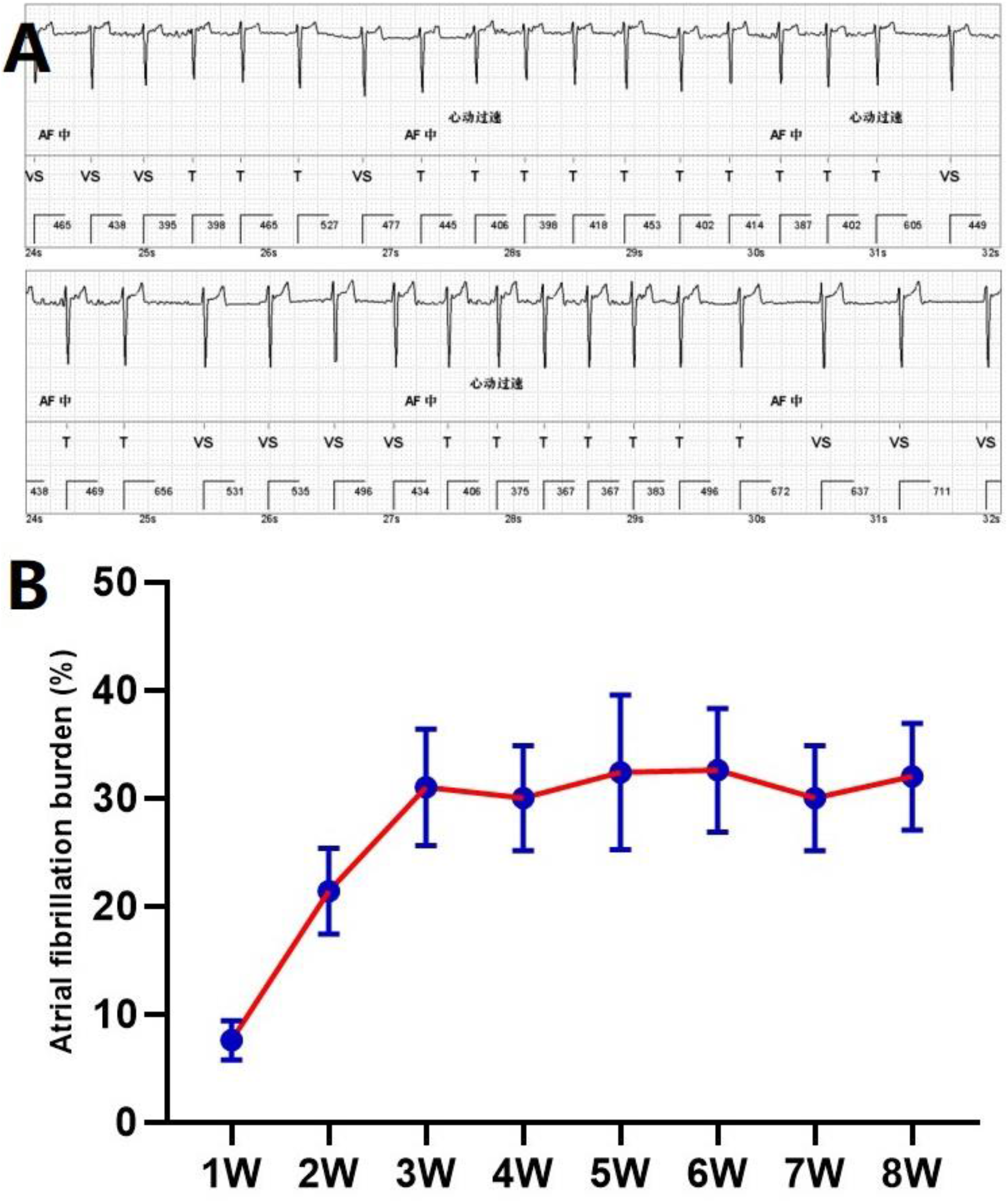
Recorded atrial fibrillation and atrial fibrillation burden in the PAH group. A: atrial fibrillation recorded by insertable cardiac monitors. B: the atrial fibrillation burden in the PAH group after the injection of dehydromonocrotaline.

#### Electroanatomical mapping

for atrial myocardium, regions with locally mapped bipolar voltage amplitudes greater than 0.5mV were defined as healthy myocardium, and regions with locally mapped bipolar voltage amplitudes between 0.1 and 0.5mV were defined as low voltage tissue regions. For ventricular myocardium, the criteria for healthy and low voltage regions were local bipolar voltage amplitudes greater than 1.5 mV and between 0.5 and 1.5 mV, respectively. Set the amplitudes of local bipolar voltage in order of red, orange, yellow, green, blueness, blue, purple color gradient arrangement. Healthy myocardial tissues were shown in purple, and regions of low voltage were shown in other colors. The endocardial electroanatomical mapping showed that there were some low voltage regions in LA and RA and few low voltage regions in LV and RV in the PAH group, and the percentage of low voltage regions in the RA was larger than that in the LA (7.66±0.46% vs. 4.40±0.55%, P <0.0001). In contrast, the endocardial electroanatomical mapping in the control group did not reveal any low voltage regions. (**Figure 4**) The epicardial electroanatomical mapping of the PAH group and control group showed no obvious low voltage region.

**Figure 4.**
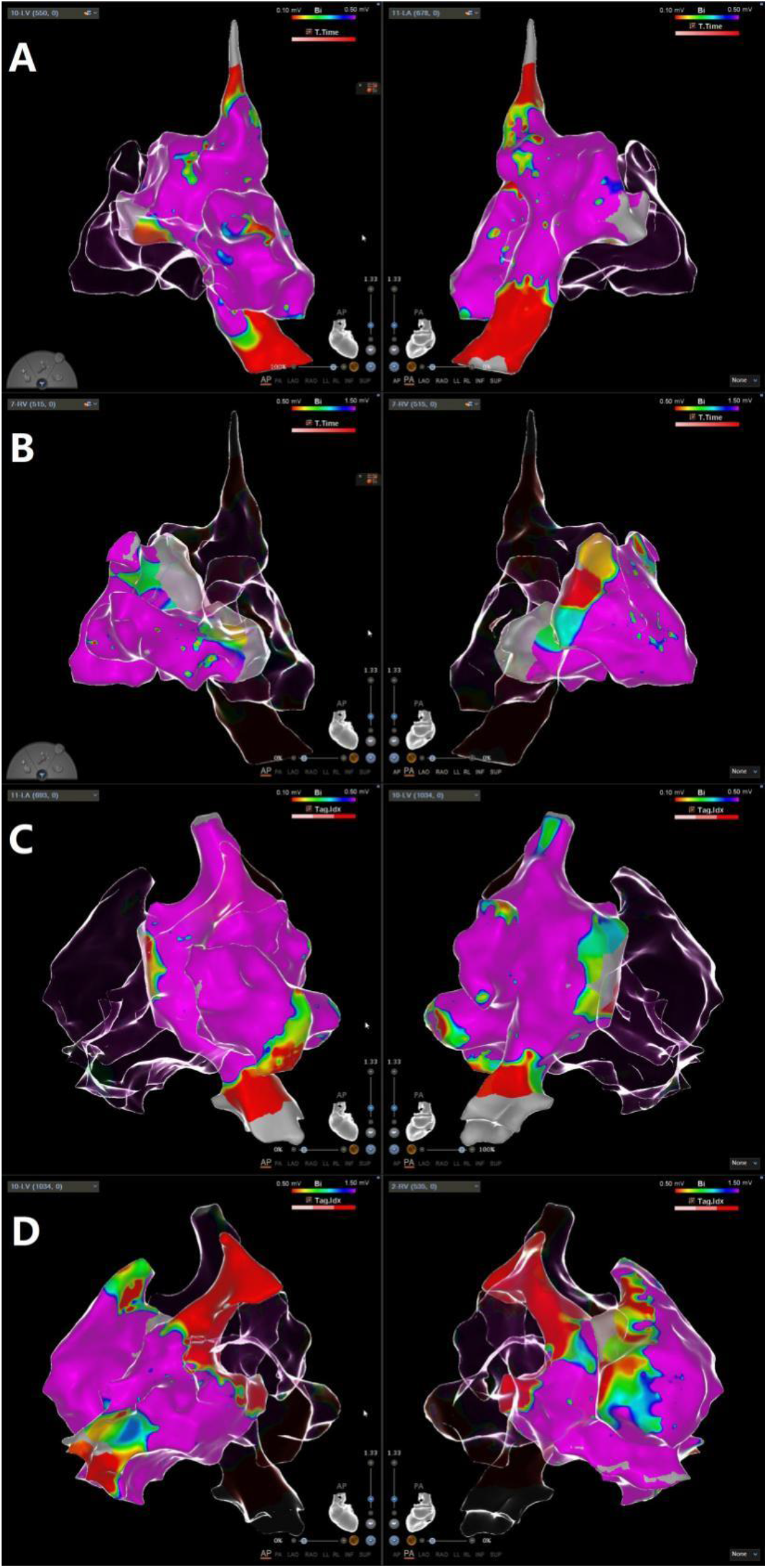
Results of endocardial electroanatomical mapping. A: atrial endocardial electroanatomical mapping in the PAH group; B: ventricular endocardial electroanatomical mapping in the PAH group; C: atrial endocardial electroanatomical mapping in the control group; D: ventricular endocardial electroanatomical mapping in the control group.

#### Programmed stimulation

the results of programmed stimulation showed that the ERPs of atriums and ventricles in the PAH group were shorter than those in the control group (RA: 128.2±5.8ms vs. 178.0±6.0ms, P < 0.0001; LA: 134.4±4.0ms vs. 174.6±5.7ms, P < 0.0001; RV: 222.6±12.4ms vs. 263.6±6.7ms, P=0.001; LV: 218.8±10.8ms vs. 259.8±5.3ms, P <0.0001). Atrial dERPs in the PAH group were significantly higher than those in the control group (RA: 0.27±0.04 vs. 0.21±0.02, P=0.024; LA: 0.29±0.02 vs. 0.25±0.03, P=0.022), while there was no significant differences in ventricular dERPs between the two groups (RV: 0.20±0. 02 vs. 0.18±0.01, P=0.199; LV: 0.12±0.03 vs. 0.11±0.03, P=0.467). AF was induced in all the PAH beagles by programmed stimulation of the left and right atriums (**Figure 5**), and the RA was more likely to induce AF than the LA (WOV: 39.0±6.5ms vs. 28.0±5.7ms, P=0.022). In the control group, atrial programmed stimulation could only induce two or three atrial premature beats.

**Figure 5.**
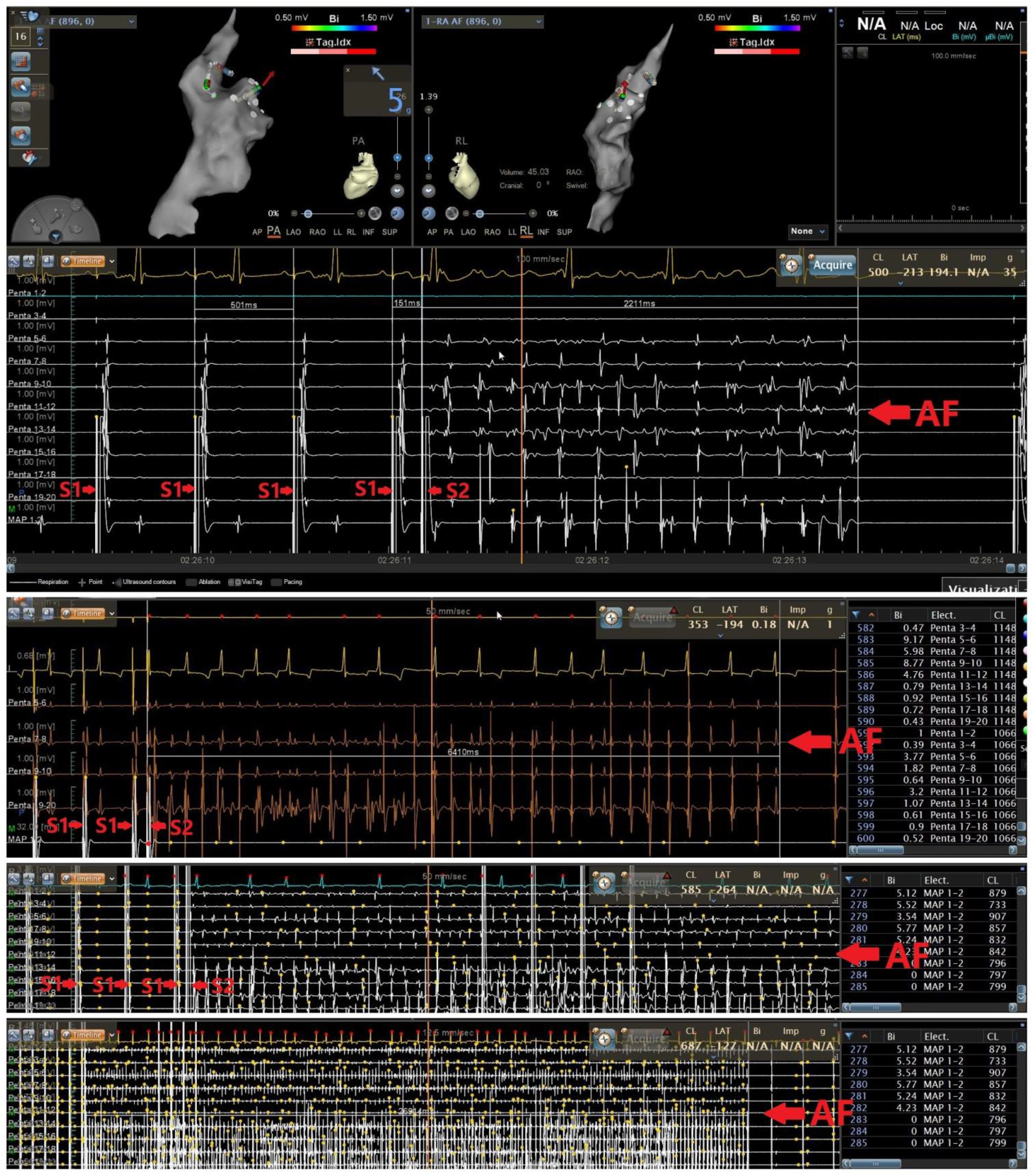
Atrial programmed stimulations could induce atrial fibrillation of varying degrees in the PAH group. AF, atrial fibrillation.

#### Myocardial conduction mapping

the conduction velocities of left and right atrial myocardium were calculated using the LAT software module and the design line function and presented by the isochronal function (**Figure 6**). The results showed that the conduction velocities of the left and right atrial myocardium of the PAH group were significantly lower than those of the control group (LA: 50.9±3.7cm/s vs. 68.2±4.1cm/s, P<0.0001; RA: 44.2±3.1cm/s vs. 67.9±3.0cm/s, P <0.0001). In addition, in the PAH group, the conduction velocity of myocardium in RA was slower than that in LA (P=0.015).

**Figure 6.**
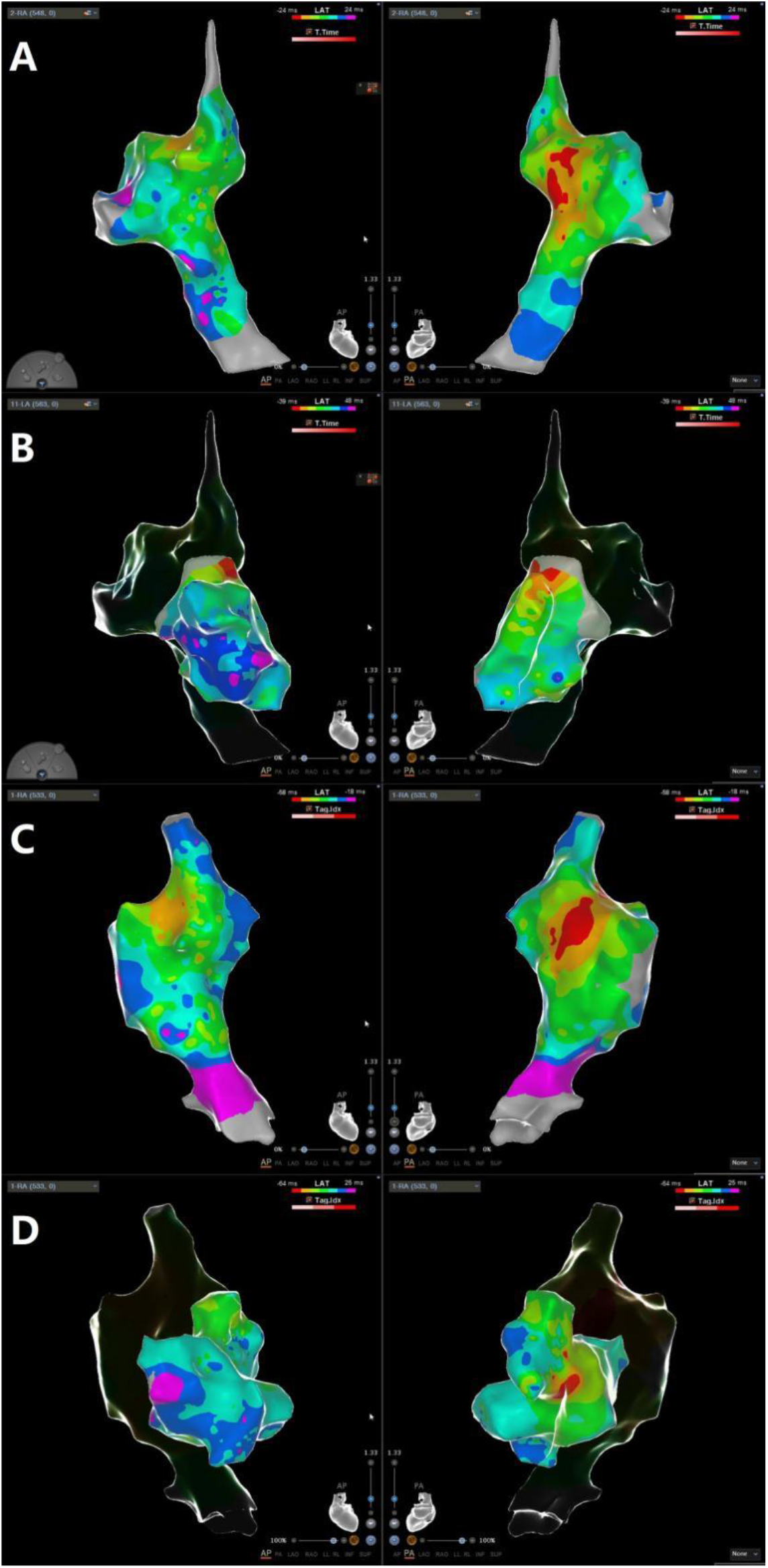
Results of myocardial conduction mapping. The more colors that appeared in a certain region, the slower the myocardial conduction velocity was. A: right atrial myocardial conduction mapping in the PAH group; B: left atrial myocardial conduction mapping in the PAH group; C: right atrial myocardial conduction mapping in the control group; D: left atrial myocardial conduction mapping in the control group.

### Histopathological studies

Masson’s trichrome staining results showed that there was a certain degree of atrial fibrosis in the PAH group, and the degree of fibrosis in RA was higher than that in LA (8.22±0.61% vs. 4.93±0.60%, P <0.0001) (**Figure 7**). In the control group, there was no obvious fibrosis in atriums.

**Figure 7.**
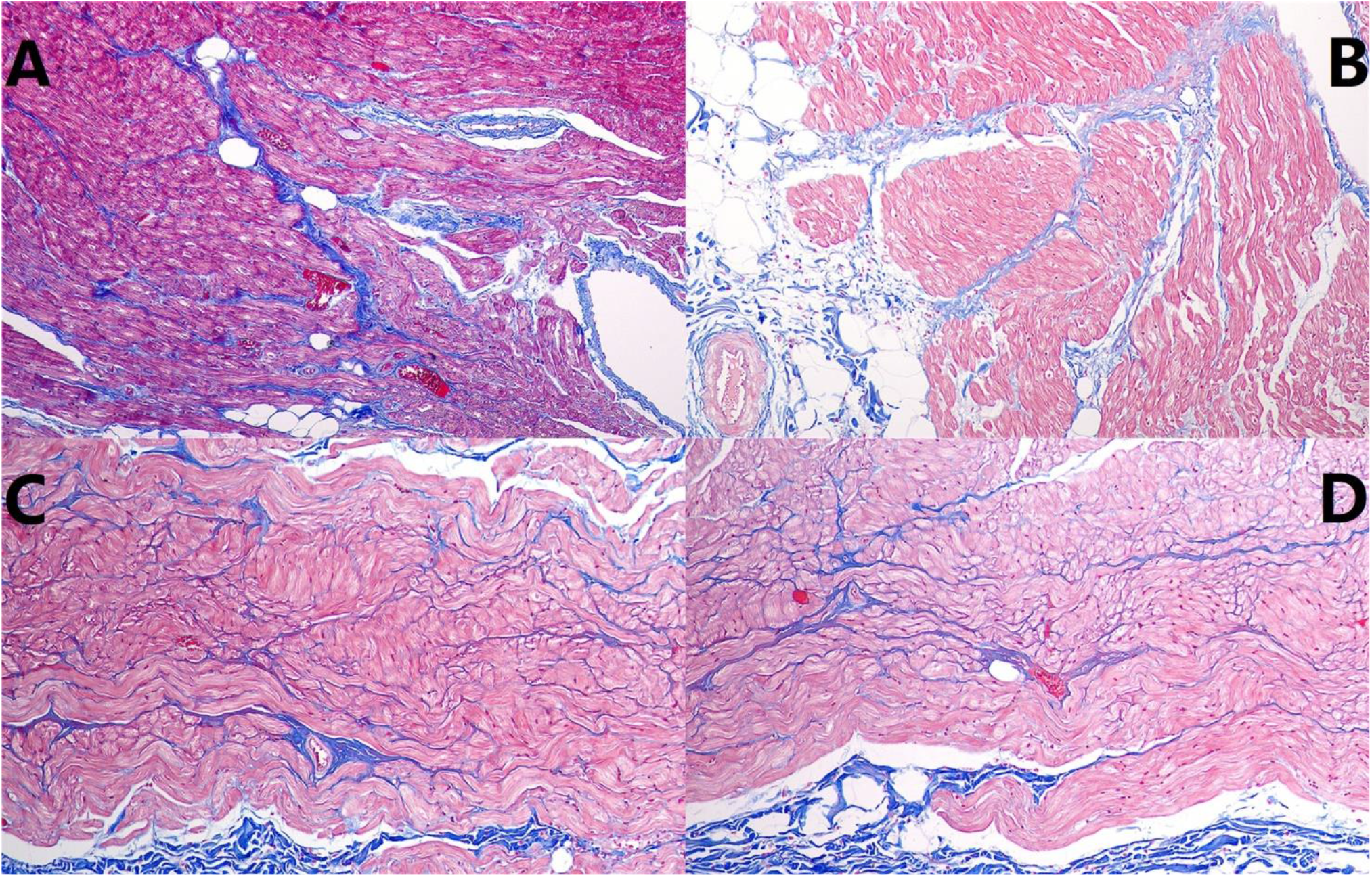
Results of Masson’s trichrome staining of right and left atrium in the PAH group (×100). A, B: right atrium; C, D: left atrium.

## Discussion

In PAH, chronic overload of the RV leads to functional tricuspid regurgitation with a subsequent increase in right atrial pressure. The longstanding elevation of atrial pressure, owing to increased pulmonary vascular resistance (PVR), induces progressive atrial dilatation and electrophysiological remodeling which, in conjunction with autonomic system modulations, may promote the development of AF.^[10]^ The increase in PVR is related to different mechanisms, including vasoconstriction (associated with smooth muscle and endothelial cell dysfunction), proliferation and obstructive remodeling of the pulmonary vascular wall, inflammation, and thrombosis. Subsequent increase in PVR leads to increased afterload in the RV, causing hypertrophy and expansion of the RV, and upstream expansion of the RA. In addition to the presence of autonomic systemic regulation present in PAH patients, these factors combine to form the basis for the initiation and maintenance of atrial arrhythmias.^[11]^ After the injection of DMCT, PAH develop over the following 2 weeks,^[12]^ consistent with a rapid increase in AF burden 2-3 weeks after the injection in this study.

There is often a causal relationship between the development of AF and the clinical deterioration in patients with PAH. This clinical deterioration seems to be associated with loss of ventricular transport mechanisms and/or detrimental hemodynamic effects of impairment of diastolic filling time due to rapid heart rate in the presence of ventricular dysfunction. As the development of AF can cause significant hemodynamic instability,^[13]^ AF may be a particularly malignant arrhythmia in patients with chronic PAH and right heart failure.

### Electrophysiological mechanisms of PAH leading to AF

Long-term PAH causes elevated atrial pressure and increases the propensity to develop AF by changing the electrophysiological properties of the atrial tissue.^[10]^ Chronic atrial stretching is the basis of a range of clinical conditions leading to arrhythmia, and this stretching of atrial myocardium could come after a variety of diseases, such as hypertension,^[14]^ heart failure,^[10]^ valvular heart disease,^[15]^ and so on.

Although animal models and clinical experience suggest a potential association between PAH and AF, little is known about how PAH leads to AF. In this well-established beagle DMCT-induced PAH model, we found that: 1) the inducibility of AF was greatly enhanced, especially in the RA;2) the atrial ERP of the PAH group was significantly shortened and its dERP was significantly increased; 3) the conduction velocities of the atrial myocardium decreased significantly, and the decrease of RA was more obvious. That was, the atria, especially the RA, produced significant electrophysiological remodeling, which provided the electrophysiological basis for the onset of AF.

It was reporteded that, in population with AF and chronic lung diseases with elevated mPAP, non-PV arrhythmogenic foci were localized in RA in as much as 26.7% of patients.^[16]^ Therefore, in addition to conventional left atrial ablation, more attention should be paid to the mapping and intervention of the RA during catheter ablation for AF patients with PAH.

### Hypoxia and AF

In this study, PAP in the PAH group was not as high as reported in previous literature,^[17]^ which may be related to the significantly larger size of beagles used in this study. However, the incidence and inducibility of AF in PAH beagles in this study increased significantly. Following exposure to the DMCT, the lung begins to have a series of changes, such as pulmonary vascular remodeling,^[18]^ increased lung mass, and microthrombi formation,^[19,20]^ which are common in diseases that can lead to PAH. Thus, we make a point: PAH begins to lead to an increased susceptibility to AF before it becomes severe, and this may be associated with hypoxia.

Nocturnal hypoxemia is an independent risk factor for AF.^[21]^ Hypoxia could lead to a significant increase in conduction inhomogeneity and decrease conduction velocity, both leading to atrial arrhythmias.^[22]^ Hypoxemia can also cause increased vagal nerve tension, which aggravates bradycardia and conduction rhythm disorders in hypoxic patients.^[23]^ Thus, the oxidative stress response to hypoxia can promote electrical remodeling and increase the incidence of arrhythmia. In addition, sympathetic upregulation due to hypoxemia and the presence of chronic systemic inflammation may also contribute to the development of AF.^[24]^

### Comparisons with other animal models of AF

As human AF is heterogeneous, multifactorial, and slowly developing, animal models may only mimic particular aspects of human AF without representing the complex pathophysiology in its entirety.^[25]^ Therefore, a variety of animal models of AF have been developed for different causes.

Because of their accessibility and the ease with which they can be generated and manipulated, rodents have been used extensively to study AF. Rodents can be applied to make many models of AF through genetic reprogramming^[26,27]^ and simulation of multiple clinical risk factors.^[28-30]^ Similar to this study, Hiram R et al^[8]^ studied the right atrial mechanisms of AF in a rat model of right heart disease. However, their rat model of AF required stimulation to induce AF, while our model could present with spontaneous AF. In addition, because the hearts of rats were too small, they could not use tools that could be used for human electrophysiology as in this study.

Dogs have been widely used to study the influence of inflammation,^[31]^ autonomic nervous system^[32]^ and congestive heart failure on AF.^[33]^ Shi et al has studied atrial remodeling secondary to ventricular remodeling through a model of rapid ventricular pacing in canines.^[34]^ Although the right cardiac system was intervened, their model did not reflect the difference in AF induction between the left and right atria as our model did. The model in this study was also more similar to the real clinical situation.

For other big animals, due to their relatively good tolerance to AF, goats can be used to construct a model of persistent AF, which is beneficial to study the structural and electrophysiological remodeling in AF.^[35,36]^ The goat model is usually constructed by pacing, which is very different from the pathophysiological mechanism of AF in our model.

## Conclusion

DMCT-induced PAH can cause stable spontaneous AF and high AF inducibility in cannine model. The exact mechanism is related to RA electrophysiological and structural remodelling.

## Acknowledgements

This research was supported by Grants from the Nature Science Foundation of China (NO. 81770324 and NO. 81670305).

